# An optimized 16S rRNA sequencing protocol for vaginal microbiome to avoid biased abundance estimation

**DOI:** 10.1101/857052

**Authors:** Qiongqiong Zhang, Lei Zhang, Ying Wang, Meng Zhao, Rui Chen, Zhi Tao, Tao Lyu, Zhenyu Huang, Qinping Liao

## Abstract

We applied three 16S rRNA sequencing protocols on vaginal microbiome samples, to evaluate whether they produce unbiased estimation of vaginal microbiome composition. We modified the 27F primer (hereafter denoted as 27F’). Using vaginal samples from 28 healthy women and 10 women with bacterial vaginosis, we sequenced three 16S rRNA sequencing protocols, i.e., 27F-338R, 27F’-338R and 341F-806R protocols, naming after their PCR primer sets, to test whether the sequencing results are consistent with the clinical diagnostics, morphology and qPCR results. First, the 27F primer would not align with *Gardnerlla vaginalis* very well, leading to poor amplification of such species. By modifying the primer sequences, the modified 27F primer (27F’) was able to amplify *Gardnerlla vaginalis* very well. Second, the DNA sequence of characteristic species *Lactobacillus crispatus* is identical with *Lactobacillus garrinarum*, leading to biased estimation of abundance of *Lactobacillus crispatus* when using V3-V4 as PCR target region; in contrast, such bias did not occur when using V1-V2 as a target region. Third, optimized 27F’-338R avoided above-mentioned biases and restored the well-established community state types (CSTs) clustering.

**Importance:** Vaginal microbiome has profound effects on the health of women and their newborns. Our study found that two well-established 16S rDNA sequencing protocols led to systementical biased estimation of characteristic species of vaginal microbiome. Subsequent analysis proved that the PCR primer fetching efficacy and target region identity were major contributor for such bias. With carefully selected target region and optimized PCR primer set, we were able to eliminate such biases and provide accurate estimation of vaginal microbiome, which showed high consistency with clinical diagnostics. We modified the 27F primer (27F’). Using the optimized PCR primer set of 27F’ and 338R to target the V1-V2 hyper-variable region, our 16S rRNA sequencing correctly evaluate the composition of vaginal microbiome.

## Introduction

The vaginal microbiome has been recognized as a critical factor involved in the protection of the female from various bacterial, fungal and viral pathogens.(1) Bacterial vaginosis (BV) is the most common lower reproductive tract infectious disease in reproductive age women. It is associated with a range of health issues such as pelvic inflammatory disease,(2–4) infertility,(5) preterm delivery,(6) tumors(7, 8) and sexually transmitted diseases.(9–11) Vaginitis was previously diagnosed by culturing bacteria in the vagina, which may overlook some fastidious bacteria that have not been isolated by culture.(12) Nowadays, the diagnosis of BV is typically made by Amsel criteria(13) or Nugent score.(14)

With the advent of high-throughput sequencing methods, more and more studies have proposed 16S rRNA sequencing to estimate the composition of vaginal microbiome.(15–17) Partial amplification of bacterial 16S gene sequences with primers across hypervariable regions, mainly including V1-V2 region(15, 18) and V3-V4 region,(17, 19, 20) is a common method to describe vaginal bacterial populations. However, it has been shown that different selection of primers for amplification can bias the results of 16S amplicons for microbiome studies.(21) For example, it has been reported that the universal bacterial 27F primer (5’-AGAGTTTGATCCTGGCTCAG-3’) is not suitable for targeting vaginal bacteria in BV such as *Gardnerella vaginalis*.(22) Thus the V1-V2 region primers (27F-338R) did not efficiently evaluate the microbiome in BV.(23)

Based on the above research, we modified the sequence of the 27F primer (hereafter denoted as 27F’). And we sequenced three 16S rRNA sequencing protocols, i.e., 27F’-338R, 27F-338Rand 341F-806R protocols, naming after their PCR primer sets, to test which provides the best species-level resolution of the vaginal microbiome by means of *in silico* analysis and experimental evaluation.

## Results

### 27F-338R and 341F-805R 16S rRNA protocols could not estimate female vaginal microbiome accurately

We first checked whether the widely used 27F-338R and 341F-805R 16S rRNA protocols were capable of evaluating the vaginal microbiome from women accuratly. 16S rRNA sequencing was applied on the collected vaginal swab samples from 28 healthy women and 10 women with BV. As shown in **Table 1**, the top 10 bacteria that showed highest abundance across all the samples were denoted as the representative bacteria of vaginal microbiome. For each sample, any representative bacteria with abundance over 10% was denoted as a major species (highlighted in bold and italic) and others are labeled not detected (ND).

**Table 1.**
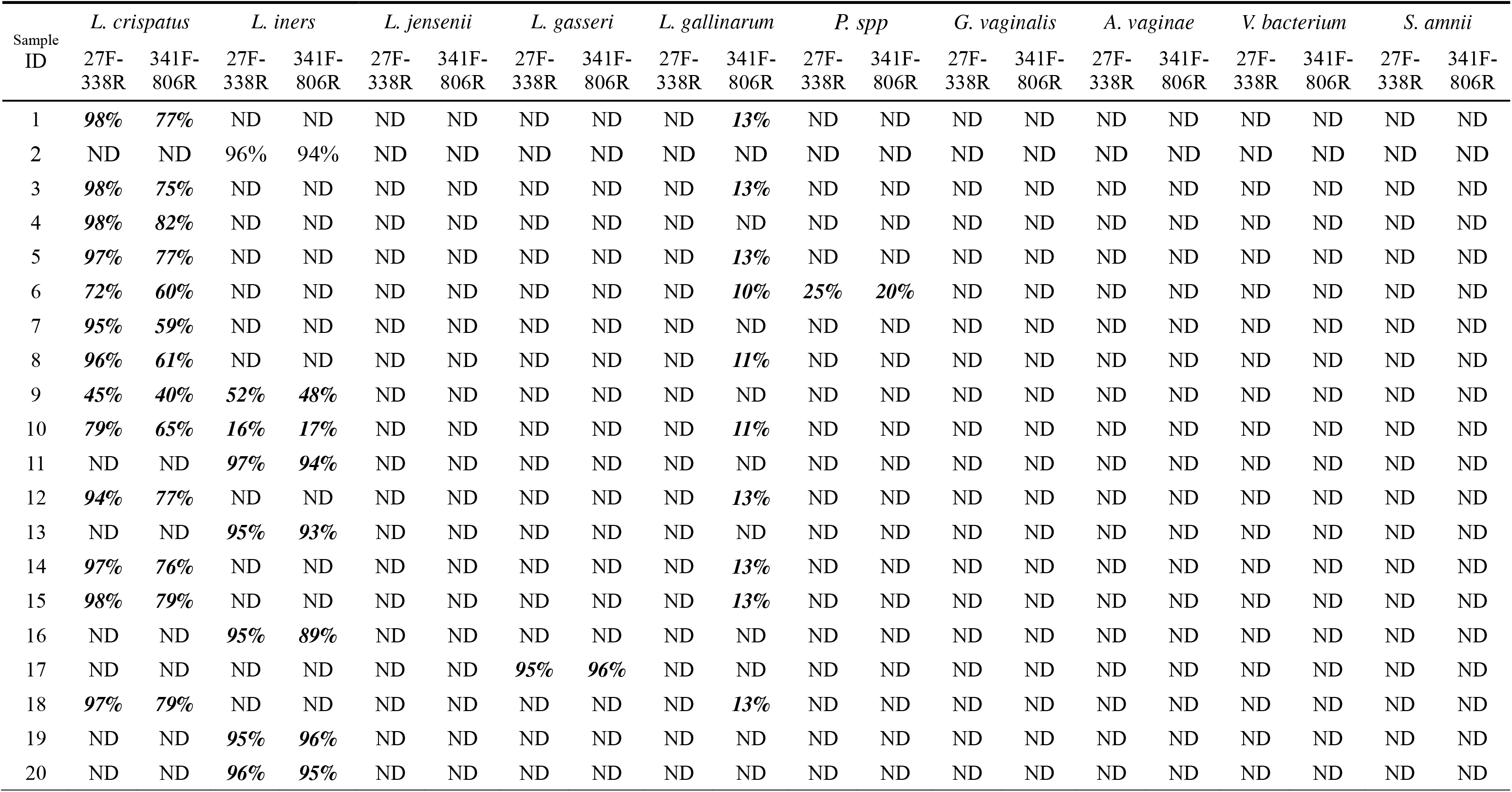
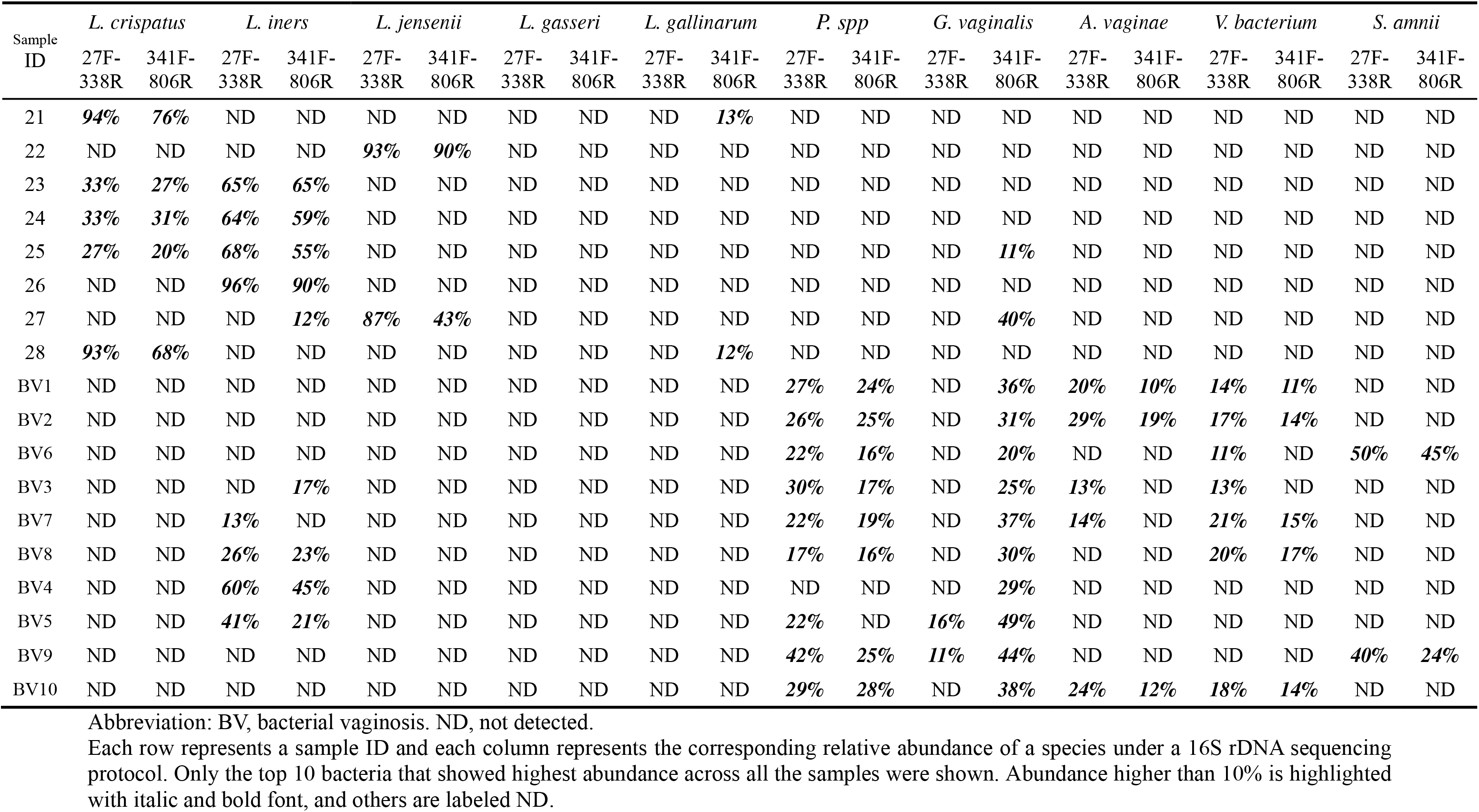
Summary of vaginal microbiome compositions from healthy and BV samples.

First, the abundance of *Gardnerella vaginalis* showed a significant difference between 27F-338R and 341F-805R protocols: in the 27F-338R protocol, only 2 out of 10 BV samples (20%) showed *Gardnerella vaginalis* as a major species, while in 341F-805R protocol, 10 out of 10 BV samples (100%) showed *Gardnerella vaginalis*. *Gardnerella vaginalis* was confirmed by morphology and microscope results in all the BV samples (**Appendix Figure 1**), thus the 341F-805R protocol is more accurate in women. What’s more, with *Lactobacilli* and *Gardnerella vaginalis* specific primers, our qPCR validation from 15 random samples also supported the results of 341F-805R protocol (**Appendix Figure 2**).

It was also noted that another unexpected bacterium, *Lactobacillus gallinarum*, showed up as a major species in 12 out of 28 healthy samples (43%) from the 341F-805R protocol results. In contrast, no samples showed *Lactobacillus gallinarum* are from the 27F-338R protocol results. To our knowledge, unlike *Lactobacillus crispatus*, *Lactobacillus gasseri*, *Lactobacillus iners*, and *Lactobacillus jensenii*, *Lactobacillus gallinarum* is not a common *Lactobacilli* in vaginal microbiome.(15) We reasoned that the differences between 16S rRNA protocol may be responsible for such controversial results regarding *Gardnerella vaginalis* and *Lactobacillus gallinarum*.

### Biased abundance estimations were caused by low fetching efficacy of primer 27F and identical sequences in the V3-V4 target region

We quantified the differences between the 27F-338R and 341F-805R 16S rRNA protocols by the fetching efficacy of primer set and the identity of target regions. To do so, we evaluated the alignments of primer set and target region to the reference databases. To eliminate the potential bias caused by certain reference database, we tested two databases in parallel, i.e., SLIVA and NCBI 16S Microbioal database.

First, we aligned the PCR primer sequences of 27F, 338R, 341F and 805R to the reference 16S rRNA sequence databases to evaluate the primer fetching efficacy. As shown in **Figure 1A**, 27F primer could not align all of the reference sequences (88.9% in SLIVA database and 57.3% in NCBI 16S Microbioal database), compared to 100% for 338R, 341F and 805R primers (in both databases). Two species, i.e., *Gardnerella vaginalis* and *Bifidobacterium bifidum*, were found unable to align with the 27F primer. Another human vaginal microbiome characteristic species, *Atopobium vaginae*, was also found imperfect match with the 27F primer. This is consistent with a previous work that argued 27F primer could reduce PCR efficiency.(22) This also explained why the *Gardnerella vaginalis* was negligible in low abundance from the 27F-338R protocol results.

**Figure 1:**
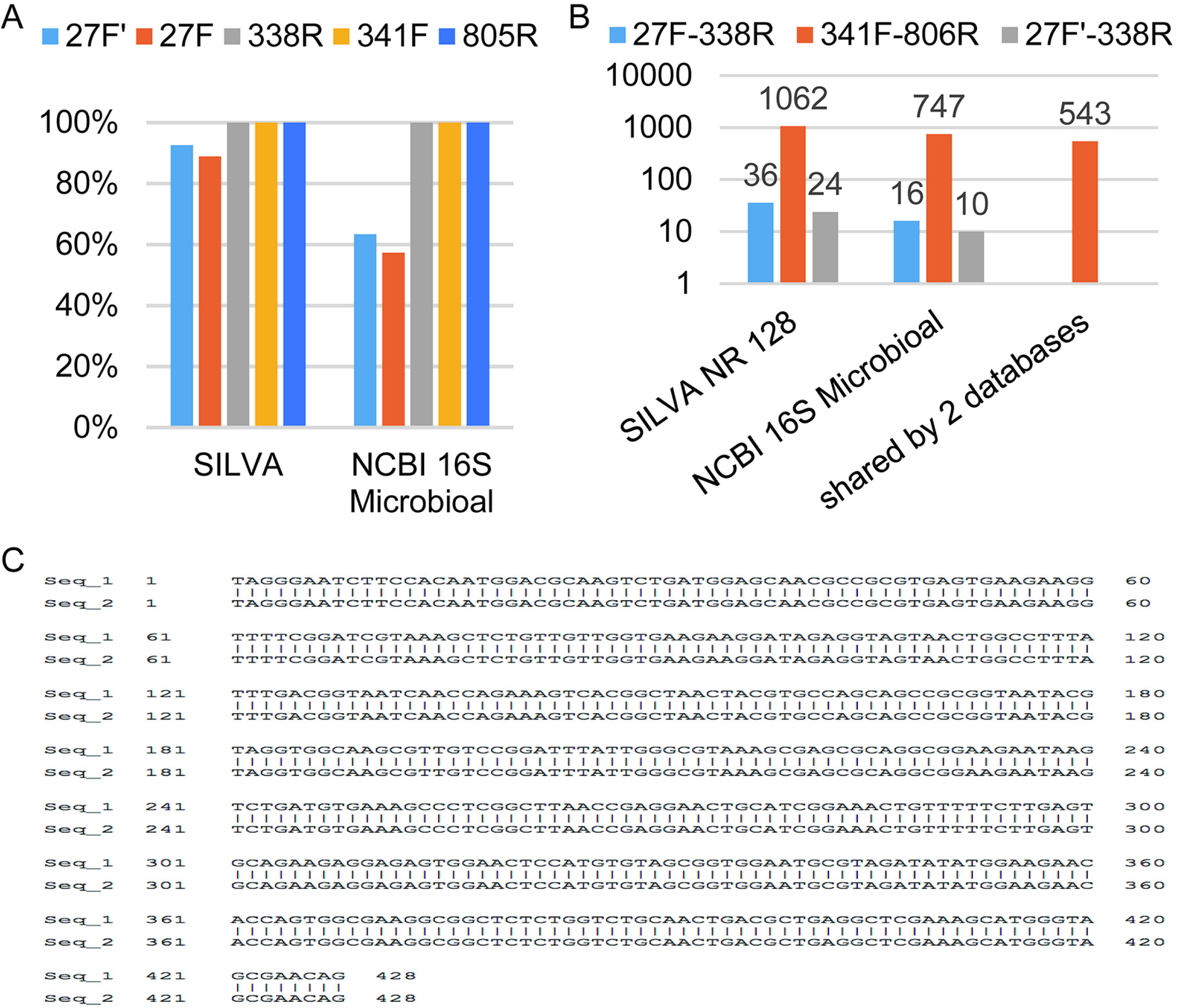
PCR primer fetching efficacy and target region identity quantification. A. Primer efficiency were quantified by the alignment of primer sequence to the reference sequences. In X-axis, two reference databases were used, SLIVA and NCBI 16S Microbioal. The Y-axis showed the percentage of aligned reference sequences by certain primer sequences, including 27F’ (blue), 27F (orange), 338R (grey), 341F (yellow) and 805R (dark blue). B. Number of identical sequences shared by two different species had been shown in bar plot. The X-axis represents the reference database we used. C. Alignment of *Lactobacillus crispatus* and *Lactobacillus gallinarum* at V3-V4 region.

Second, we extracted the target regions corresponding to primer sets of 27F-338R and 341F-805R (V1-V2 and V3-V4, correspondingly) and count the identical sequences shared by different species. As shown in **Figure 1B**, there were much more species that share identical sequences with others in the target region of 341F-805R protocol (1062 for SLIVA database, 747 for NCBI 16S Microbioal database and 543 for intersection of the two databases) than 27F-338R protocol (36 for SLIVA database, 16 for NCBI 16S Microbioal database and 0 for intersection of the two databases). We further checked the species that share identical sequences with others, and found that *Lactobacillus crispatus* share identical sequence with *Lactobacillus gallinarum*, in the target region of 341F-805R primer set (**Figure 1C**). This explained why *Lactobacillus gallinarum* showed in high abundance from the 341F-806R protocol results.

To optimize the 16S rRNA protocol, we modified the sequence of 27F primer (see **Methods** for details), to allow higher PCR fetching efficacy. The modified 27F primer was denoted as 27F’ and the corresponding 16S protocol was named as 27F’-338R protocol. As shown in **Figure 1A**, in the SLIVA and NCBI 16S Microbioal databases, the 27F’ primer aligned 92.6% and 63.4% of reference 16S rRNA sequences, correspondingly; higher than the alignment rate of 27F (88.9% and 57.3%, correspondingly). What’s more, the 27F’ primer showed perfect match with *Gardnerella vaginalis*, *Bifidobacterium bifidum* and *Atopobium vaginae*. In addition, as shown in **Figure 1B**, 27F’-338R protocol showed 24, −10 and 0 species that share identical sequences with others in the target region, from reference database of SLIVA, NCBI 16S Microbioal database and intersection of the two databases, correspondingly. These results indicating that our optimized 27F’-338R 16S rRNA protocol could be a better choice for human vaginal microbiome.

### Optimized 27F’-338R 16S rRNA protocol provided unbiased estimation of vaginal microbiome

We furthur validated the 27F’-338R protocol. First, we merged all the BV samples to count the abundance of the top ten bacteria for three 16S protocols (**Figure 2A**). The top 10 species found in BV condition included *Gardnerella vaginalis*, *Prevotella* spp., *Lactobacillus iners*, *Veillonellaceae bacterium*, *Sneathia amnii*, *Clostridiales bacterium*, *Atopobium vaginae*, *Chlamydia trachomatis*, *Sneathia sanguinegens and Candidatus saccharibacteria*. Overall, we noticed that the results from 27F’-338R and 341F-806R protocols were quite similar and the 27F-338R protocol seemed quite different. The *Gardnerella vaginalis*’s relative abundance is about 41%, 33% and 8%, when applying the 27F’-338R and 341F-806R and 27F-338R protocols, respectively. This indicated that the low *Gardnerella vaginalis* estimation from 27F-338R protocol was recalibrated by the 27F’-338R protocol. Second, we merged all the healthy samples to count the abundance of top bacteria under different protocols (**Figure 2B**). Unlike the BV group, the top species were mainly *Lactobacilli*, i.e., *Lactobacillus crispatus*, *Lactobacillus iners*, *Lactobacillus jensenii*, *Lactobacillus gasseri*, *Lactobacillus gallinarum*, *Gardnerella vaginalis*, *Prevotella* spp., *Lactobacillus helveticus*, *Lactobacillus acidophilus and Streptococcus anginosus*. At this time, we noticed that the 27F’-338R and 27F-338R protocols were quite similar and the 341F-806R protocol seemed quite different from others. The emerging of in-relevant *Lactobacillus* spp., i.e, *Lactobacillus gallinarum*, *Lactobacillus helveticus* and *Lactobacillus acidophilus* in the 341F-806 protocol is because of misalignment due to the identical sequence in the target region. In conclusion, we showed that the 27F’-338R protocol could recalibrate the biased estimation of *Gardnerella vaginalis* and *Lactobacillus crisptus*.

**Figure 2:**
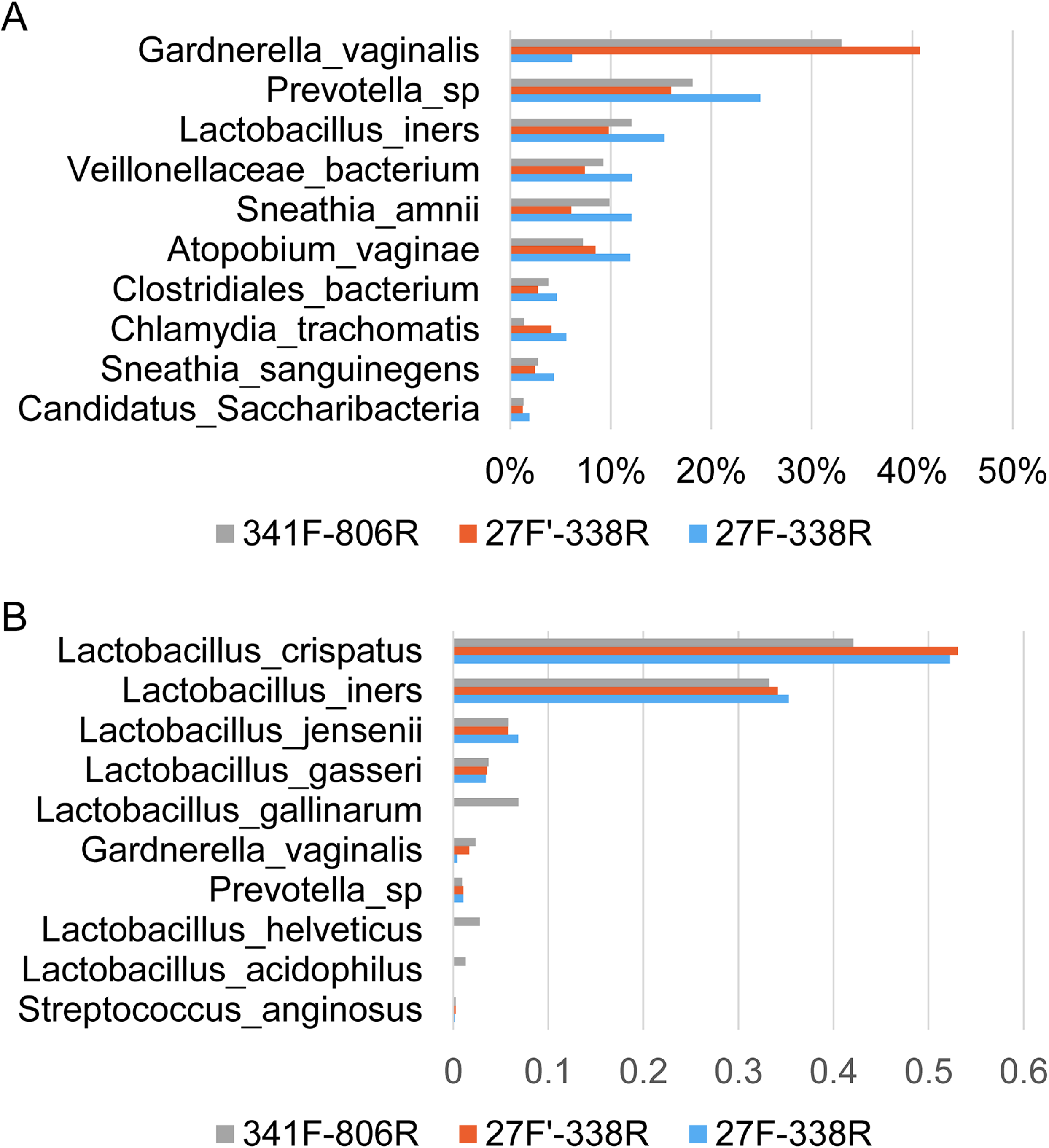
Comparison of 16S rRNA sequencing results from 27F-338R, 27F’-338R and 341F-806R protocols. A.The top ten bacteria’s abundance were from the BV group. Three protocols were compared, i.e., 27F-338R (blue), 27F’-338R (orange) and 341F-806R (grey). B. Like in A, the top ten bacteria showed in the healthy group from three protocols, i.e., 27F-338R (blue), 27F’-338R (orange) and 341F-806R (grey), were compared.

Subsequently, we found the 27F’-338R protocol could restore the well-established community state types (CSTs) clustering.(15) We performed unsupervised clustering of 28 healthy and 10 BV samples using the abundance of the top 20 bacteria (**Figure 3**). We noticed all the healthy samples were clustered together and all the BV samples were clustered together. All the BV samples showed *Lactobacillus* diminished and *Gardnerella vaginalis* dominated diverse community, similar to the CST-IV cluster.(15) For the healthy samples, we noticed all *Lactobacillus crispatus* enriched samples were clustered together, so were the *Lactobacillus gasseri* enriched samples, the *Lactobacillus iners* enriched samples and the *Lactobacillus iners* enriched samples; and they formed the CST-I, CST-II, CST-III and CST-V cluster.(15) In summary, we propose that the 27F’-338R protocol based 16S rRNA sequencing method could give an unbiased estimation of vaginal microbiome.

**Figure 3:**
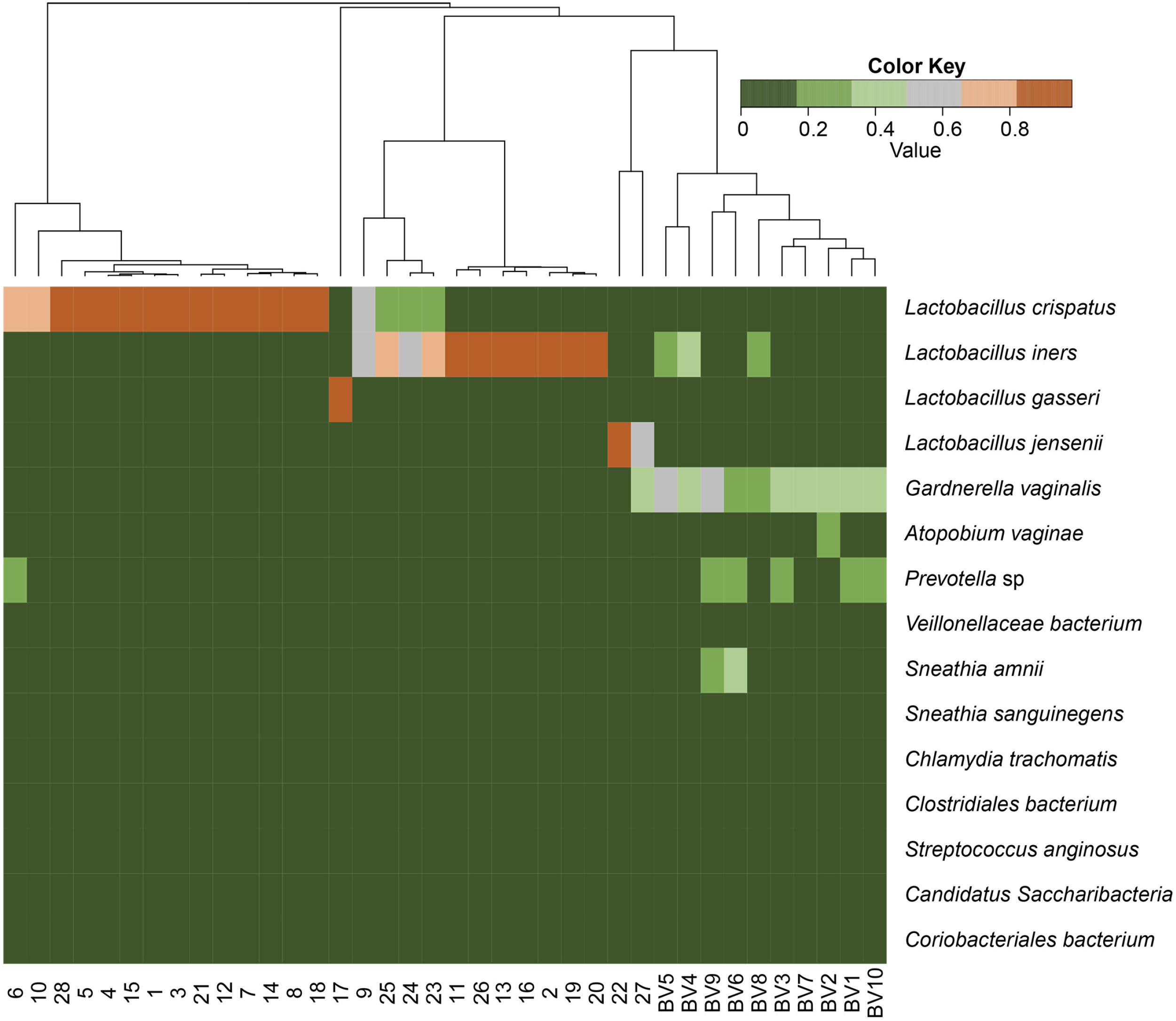
Heatmap and dendrogram of vaginal compositions from 28 healthy and 10 BV samples. The vaginal compositions from 28 healthy and 10 BV samples utilizing 27F’-338R protocol were clustered and colored by relative abundance (from low to high abundance, color changes from green to red).

## Disscussion

16S rRNA sequencing has been used to identify the bacterial composition of the human vaginal microbiome in multiple ethnic groups, but the study on the population’s vaginal microbiome is still insufficient. In addition, no studies have examined whether different 16S rRNA sequencing protocols are an unbiased way to identify vaginal microbes. Our principal findings were that the 27F primer was not well aligned with *Gardnerlla vaginalis*, resulting in poor amplification effect. By modifying the 27F primer, 27F’ could well amplify *Gardnerlla vaginalis*; The DNA sequence of *Lactobacillus crispatus* was the same as that of *Lactobacillus garrinarum*. There was a bias in the estimation of *Lactobacillus crispatus* abundance when V3-V4 was the target region of PCR, while there was no such bias when V1-V2 was the target region; The optimized 27F ’-338R avoids the above deviation and restores the well-established community state types (CSTs) clustering.

As we showed in the introduction section, a series of 16S rRNA sequencing protocols with different target regions and corresponded primer sets were utilized in vaginal microbiome studies. However, due to the limit on reads length, only a subset of target regions remains available. One recent study had performed in-silico and experimental evalutions on primer sets of V1-V3, V3-V4 and V4. In their conclusion, V4 region provides the best results on species level resolution of the vaginal microbiome.(21) In our evaluation, we emphasized the consistency between the 16S rRNA sequencing results and clinical diagnosis, such as morphology and culture of the characteristic species. Another study compared two 16S rRNA protocols, utilizing V1-V2 and V3-V4 hypervariable regions as target regions. They found 16S rRNA sequencing protocol utilizing V3-V4 hypervariable region would identified more species and the ones using V1-V2 hypervariable region would miss several characteristic speices of vaginal microbiome.(23) We agreed with them that unoptimized 16S rRNA sequencing protocol utilizing V1-V2 hypervariable region would produce biased estimation.

*Gardnerella vaginalis* is a well recognized bacteria, which is confirmed by morphology and microscope results in all the BV samples. However, through our *in-silico* analysis, *Gardnerella vaginalis* were found unable to align with the 27F primer. This is consistent with previous reports as the 27F primer could not match the *Gardnerella vaginalis* very well, leading to a low PCR efficiency. ^22^ For other microbiome, if we normalized the *Gardnerella vaginalis*’s abundance, they showed no significant difference under the 27F’-338R and 341F-806R and 27F-338R protocols.

*Lactobacillus* spp. are so important in human vaginal microbiome that four *Lactobacillus* spp. were the characteristic species used by the authoritative five community state types (CSTs), which are established to group vaginal microbiome patterns according to the dominant species present: CSTI, II, III, IV and V dominated by *L.crispatus*, *L. gasseri*, *L. iners*, diverse community and *L. jensenii*, respectively.(15) However, we found that *Lactobacillus crispatus* share identical sequence with *Lactobacillus gallinarum* when using the target region of 341F-805R primer set. That is, if we used the V3-V4 as the target region, we might wrongly assign the characteristic species of CST-I (*Lactobacillus crispatus*) to another vaginal microbiome in-relevant species (*Lactobacillus gallinarum*).

As shown in our trial experiments, the 27F-338R protocol under-estimated the abundance of *Gardnerella vaginalis*. In addition, we showed that 16S rRNA sequencing protocol utilizing V3-V4 hypervariable region would also introduce bias: the 341F-806R protocol misaligned *Lactobacillus crisptus* to other in-relevant *Lactobacilli*. What’s more, these biases only occurs in its own protocol, but could not be repeated in the other protocol. Therefore, we reasoned that such bias was not sample or ethnic group related, but instead, associated with unoptimized 16S rRNA sequencing protocols. We have pinned down that primer sequence and target region are the major contributor for the bias. Subsequently, we have optimized the protocol, using the modified 27F primer and chose the V1-V2 hyper-variable region as the target region. The optimized 16S rRNA sequencing protocol had been proven to be able to recalibrate the estimation of *Gardnerella vaginalis*, preventing misalignment of *Lactobacillus crispatus* and restored the authoritative five community state types (CSTs).

This study provides an optimized 16S rRNA-based protocol for evaluating the composition of human vaginal microbiome using current common NGS sequencing platform. and it is the first piece of work that systematically investigated the female vaginal microbiome with above-mentioned methods. This optimized 16S rRNA-based protocol can not only accurately assess the composition of vaginal flora, but also accurately and economically. The accurate assessment of vaginal microbiome could contribute to the treatment of vaginitis in hospital.

Serval further works will be updated regard the following aspects. In this study, we used BV sample and healthy samples, because the vaginal microbiome is mainly dominated by bacteria in these two groups. Another bacterium dominate disease, aerobic vaginitis, will be tested in our subsequent work. Yet, one disadvantage of the 16S rRNA sequencing was exposed, as well and that is that the 16S rRNA sequencing is not suitable for the diagnosis of TV, VVC, HPV, HIV and so on. Currently, we used the clinical diagnostics such as such as morphology and culture of the characteristic species as ground truth of human vaginal microbiome’s composition. However, the composition of human vaginal microbiome is constantly being updated as more and more new technologies are being applied, such as metagenome related technology. It should also be noted, that as we were restricted by the sequencing platform, we only tested the target regions of the V1-V2 and V3-V4, leaving the V1-V3, V4, V4-V6 target regions unexamined, albeit future work will examine such target regions not included in the present study.

## Materials and methods

### 27F’ primer design

As mentioned above, the common nondegenerate form of the 27F primer (5’-AG**A**GTT**T**GAT**C**CTGGCTCAG-3’) is not suitable for targeting *Gardnerella vaginalis* in BV.(22) Meanwhile, the sequence (5’-AG**G**GTT**C**GAT**T**CTGGCTCAG-3’) most frequently observed binding site sequence is found in *Bifidobacteriales*, including the genus *Gardnerella* (GenBank accession numbers M58729 to M58744).(22, 24) Its binding site variant is of particular interest to the study of vaginal microbiology in BV, and the sequence has three mismatched bases compared to the common sequence of the 27F primer. To combine two sequences’ strengths, we merged their different bases (R=A/G,Y=T/C), and got an modified 27F primer, i.e., 27F’ (5’-AG**R**GTT**Y**GAT**Y**CTGGCTCAG-3’).

### Study Population

28 healthy women without vaginitis such as aerobic vaginitis (AV), bacterial vaginosis (BV), vulvovaginal candidiasis (VVC), and trichomonas vaginitis (TV), and 10 women with BV only were enrolled at the gynecological clinic of Beijing Tsinghua Changgung Hospital from April to October 2018. All women were aging between 18 and 50 years old and were not pregnant or breast-feeding. The protocol was approved by the Medical Ethics Committee of Beijing Tsinghua Changgung Hospital. Written informed consents were obtained from each participant.

### Sample collection and DNA Extraction

The vaginal secretions were obtained via two swabs. One swab was used to prepare a dry slide for Gram staining, under 400× magnification for visual detection, to test for AV, BV, VVC, and TV. The criteria of Donders(25) et al. was used to diagnose AV (with a score of 3 or greater). BV was determined by Nugent’s criteria (Nugent score of 7 or greater).(14) The diagnosis of VVC and TV was mainly based on morphological observation under high power field (400× magnification). The other swab was quickly plunged into a tube containing 1 ml PBS solution and stored at −80°**C** until total DNA extraction of vaginal flora. The DNA of the sample was extracted through the TIANamp Bacteria DNA Kit (TIANGEN, China) according to the manufacturer’s instructions. This step required additional Lysozyme (Sigma–Aldrich), proteinase K, RNase A (Sigma–Aldrich), and finally washed and stored the DNA with 1×TE buffer. A spectrophotometer was used (Thermo Scientific NanoDrop One) to measure the concentration and purity of the DNA extracts. Then isolated DNA was stored at −20°**C** until needed.

### Sequencing

Taking data volume, sequencing accuracy, read length and economic factors into account, in this study, we chose the pair-end Illumina Solexa sequencing platform over 454 pyrosequencing platform. The V1-V2 and V3-V4 regions of the 16S rRNA were then separately amplified with universal primers 27F (5’-AGAGTTTGATCCTGGCTCAG-3’) and 338R (5’-GCTGCCTCCCGTAGGAGT-3’), 341F (5’-CCTAYGGGRBGCASCAG-3’) and 806R (5’-GGACTACNNGGGTATCTAAT-3’). The V1-V2 regions were also amplified with our modified primers 27F’ (5’-AGRGTTYGATYCTGGCTCAG-3’) and 338R (5’-GCTGCCTCCCGTAGGAGT-3’). All PCR reactions were carried out with Phusion® High-Fidelity PCR MasterMix (New England Biolabs). The PCR products examined with 400-450bp were chosen and mixed in equal density ratios. Then, the mixture PCR product was purified with Qiagen Gel Extraction Kit (Qiagen, Germany). Sequencing libraries were generated using a TruSeq® DNA PCR-Free Sample Preparation Kit (Illumina, USA) following the manufacturer’s recommendations and index codes were added. The library quality was assessed on the Qubit@ 2.0Fluorometer (Thermo Scientific) and Agilent Bioanalyzer 2100 system. At last, the library was sequenced on an Illumina HiSeq 2500 platform and 250 bp paired-end reads were generated.

### Reference Database

We compared SLIVA and NCBI in the following evaluations, as the Green genes database has not been updated since 2013(26) and RDP database is semi-automatic curated.(27) For the SLIVA database, we used and downloaded the SSU 128 Ref NR 99 version from https://www.arb-silva.de. For the NCBI database, we downloaded using the blast command of blastdbcmd in June 2017. All the taxonomies are summarized into species level.

### Sequencing Data Processing

Paired-end reads were assigned to samples according to the sample-specific barcode and truncated by cutting off the barcode and primer sequence. Use the software FLASH(V1.2.7)(28) to merge paired-end reads. According to the QIIME(V1.7.0)(29) quality control process, the raw tags were mass filtered under specific filtration conditions to obtain high quality clean tags.(30)

The 16S sequence reference index was built using the command “bowtie2-build”, with default parameters. All reads were aligned against the prebuild index using bowtie2, with parameter of “bowtie2 --local”. Alignments were associated to taxonomy by a sequence-id-to-taxonomy map, provided by the reference database, using a custom Perl script. Unique reads were counted for each taxonomy and abundance was calculated for all taxonomy. Species with abundance lower than 1% or reads number less than 5 were excluded.

### qPCR validation

*Lactobacilli* and *Gardnerella vaginalis* specific qPCR primer and probe sequences were synthesized as previously described.(31) DNA was amplified using SGExcel GoldStar TaqMan qPCR Mix (Sangon Biotech) on a Bio-Rad CFX96 real-time PCR detection system.

## Acknowledgments

We would like to thank the National Natural Science Foundation of China (Grant No.81671409) and Beijing Municipal Administration of Hospitals Clinical Medicine Development of Special Funding (Grant No. XMLX201605) to support this work, and thank Dr. Hu for kind help to this article and the nurses in the outpatient department of obstetrics and gynaecology of Beijing Tsinghua Changgung Hospital for their help in the specimen collection process.

**Appendix Figure 1:** Morphology of samples under 400× magnification after gram staining. A: 28 normal samples, B: 10 BV samples.

**Appendix Figure 2:** qPCR validation of the existence of Lactobacilli and Gardnerella vaginalis.

10 vaginal microbiome samples from healthy women (highlighted in blue) and 5 from women with BV (highlighted in orange) were sampled and used to perform qPCR validation. The difference between the Cq values of *Lactobacilli* and *Gardnerella vaginalis* was used.

